# FlowSpot Enables Decentralized Phenotypic and Functional Cellular Immune Profiling from Dried Blood Spots

**DOI:** 10.64898/2026.07.20.739622

**Authors:** Rachel Caddell, Suzanne Adams, David Mushatt, Monica Vaccari, Marissa Fahlberg

**Affiliations:** Spotted Tech LLC, Bronx, New York; Section of Infectious Diseases, Deming Department of Medicine, School of Medicine Tulane University, LA; Department of Immunology and Microbiology, School of Medicine Tulane University, New Orleans, LA; Division of Immunology, Tulane National Biomedical Research Center, LA

**Author notes:** These authors contributed equally to this work (Monica Vaccari, Marissa Fahlberg).

**Keywords:** Dried blood spots, Immune cell preservation, Flow cytometry, Cellular immunophenotyping, Functional immune profiling, Decentralized clinical sampling, post-shipping recovery, CD4^+^ T lymphocytes, CD8^+^T lymphocytes, People living with HIV

## Abstract

Expanding access to cellular immune analysis is essential for decentralized clinical care, clinical trials, and population-based research. However, current flow cytometry workflows require rapid processing of fresh blood, proximity to a centralized laboratory, and cold chain logistics. Although dried blood spots (DBS) have transformed decentralized molecular diagnostics, no comparable approach has enabled robust flow cytometric analysis of immune cells.

Here, we present FlowSpot, a novel platform that enables recovery of leukocytes from DBS and preserves their immunophenotypic characteristics, allowing downstream flow cytometric analysis following ambient-temperature storage and shipment. FlowSpot recovers intact leukocytes while preserving immune cell subset frequencies with strong concordance to fresh whole blood. We demonstrate its clinical utility by enabling remote CD4⁺ T cell immunophenotyping in people living with HIV, showing high agreement with routine clinical measurements across a broad range of CD4⁺ T cell frequencies. Beyond cellular phenotyping, FlowSpot extends immune monitoring to functional profiling by enabling detection of intracellular cytokine responses, including IFNγ, IL-2, and TNFα production by CD4⁺ and CD8⁺ T cells following ex vivo PMA/ionomycin stimulation.

By overcoming a longstanding barrier to leukocyte recovery from DBS, FlowSpot extends flow cytometry beyond specialized laboratories, expanding access to cellular immune analysis for clinical care, decentralized clinical trials, and population-scale immunology.

## 1. Introduction

Blood-derived immune cell analysis is essential for clinical research, therapeutic and vaccine development, clinical diagnostics, and immune biomarker discovery (1–5). Yet, current methods depend on fresh venous blood processed in centralized laboratories, requiring rapid handling, specialized personnel, and cold-chain logistics, creating significant barriers to scalable, decentralized immune monitoring (1–10).

A major contributor to this limitation is the instability of leukocytes after blood collection. Soon after collection, blood cells rapidly decay and are either analyzed or separated as peripheral blood mononuclear cells (PBMCs) and stored in ultra-cold conditions (11).

Blood storage conditions and shipping to a specialized laboratory for analysis can significantly affect data quality (12–14), as leukocytes rapidly lose viability and undergo phenotypic changes under temperature variation, mechanical stress, and processing delays (13,15). Existing mitigation strategies, including refrigeration, cryopreservation, chemical fixation, and commercial stabilization systems, provide partial solutions but introduce important trade-offs such as limited time windows for shipping and reduced compatibility with downstream assays (16).

Together, these constraints limit routine immune monitoring for patients, particularly those requiring longitudinal testing who often face substantial travel burdens. They also restrict participation in clinical research to volunteers living within proximity to centralized trial sites. As a result, clinical trials face challenges recruiting and retaining representative populations of interest costs (9,17–20). These challenges are further amplified in resource-limited and geographically underserved regions and represent a growing operational burden for pharmaceutical development and multicenter trial execution (21).

Dried blood spots (DBS) offer a simple and scalable alternative for decentralized sampling and are widely used for molecular and serological testing, including at home testing (22–24). However, their application to cellular immune profiling has been limited by technical barriers, mainly due to the challenge of eluting phenotypically intact leukocytes adherent to the cellulose matrix (25).

Here, we present FlowSpot, a platform that enables recovery of intact leukocytes from Whatman card protein safer card for quantitative flow cytometric analysis following ambient-temperature storage and shipment. We demonstrate reproducible recovery of major immune populations, including T lymphocytes, B lymphocytes, monocytes, and neutrophils, while preserving key phenotypic and functional markers, including intracellular cytokines, without the need for cold-chain transport.

To evaluate clinical relevance, we applied FlowSpot to immune monitoring in people living with HIV (PLWH), demonstrating accurate CD4⁺ T cell measurements compared with standard clinical assays across clinically relevant ranges. Given the need for frequent CD4⁺ T cell monitoring in this population, this provides a practical framework for decentralized immune testing.

Collectively, these findings establish a pathway for cellular immune analysis from simple dried blood collection, decoupling immune monitoring from centralized laboratory infrastructure and enabling broader applications in clinical care, clinical trials, and population-scale immunology.

## 2. Material and Methods

### 2.1. Procurement of healthy donor blood and preparation of FlowSpot samples for stability analysis

Venipuncture whole blood (EDTA) from ten de-identified healthy donors was purchased from Innovative Research (Michigan, United States). Samples were shipped overnight on ice from collection sites to the processing laboratory (Albert Einstein College of Medicine Biotech Incubator, Bronx, NY). Immediately upon arrival, an aliquot from each donor was stained fresh and acquired by flow cytometry on the same day. Additional aliquots were pipetted at 25 µL per spot x 5 spots per card on Whatman 903 Protein Saver cards (Cytiva). Cards were dried overnight and either used next-day or transferred to a foil pouch (Whatman Foil-Barrier Resealable Bags, Cytiva) for processing at later time points. Samples were stored at room temperature (20-25°C) or 37°C. At the designated time points, samples were processed from one DBS card using the FlowSpot system and then stained with a cocktail of fluorescent-conjugated antibodies.

### 2.2 Study population and enrollment

The study enrolled people living with HIV receiving routine clinical care at two Tulane University-affiliated HIV clinics, East Jefferson (EJ) Hospital and Tulane Total Health Clinic (TT). Participants were approached for enrollment during scheduled clinic visits at the time of provider-ordered CD4^+^ count assessments. Eligibility criteria included age ≥18 years, English-speaking ability, attendance at a routine clinical visit, and receipt of a CD4 test as part of standard care. Exclusion criteria included unwillingness or inability to provide informed consent. No exclusions were made based on race, ethnicity, sex, socioeconomic status, or religion. Written informed consent was obtained from all participants prior to study procedures. The study was approved by the Tulane University IRB (protocol #2024-809).

### 2.3 Capillary blood collection from PLWH and sample shipping

Upon consent, participants were instructed in a supervised self-collection method involving one or two finger sticks to generate three to five dried blood spots, each consisting of approximately 25-50 µL of whole blood and applied to a Whatman 903 Protein Saver Card. Samples were anonymized using numeric study codes. After collection, DBS cards were dried at room temperature for 3-18 hours and then transferred to foil pouch. DBS were stored at room temperature for a period of 1-3 days prior to shipping via USPS. Participants received compensation via ClinCard upon completion of study procedures.

### 2.4 Clinical data collection and analysis

Clinical laboratory values, including CD4^+^ T lymphocyte count and percentages, CD4/CD8 ratio, and plasma HIV viral load, were obtained from venipuncture-based tests ordered by participants’ treating clinicians as part of standard care. These data were recorded in an access-restricted Excel spreadsheet stored on Box, a secure cloud-based platform licensed for use by Tulane University. The study coordinator anonymized and shipped samples to the processing laboratory via mail courier or standard postal service using appropriately labeled and packaged containment.

Samples were analyzed and compared against clinic-reported reference values to assess test completion rates, analytical accuracy, and potential immunologic misclassification.

### 2.5 Participants feedback

Immediately following fingerstick collection and during the same clinic visit, participants completed a brief, orally administered four-question survey designed to capture user feedback. Survey domains included challenges associated with existing CD4^+^ testing methods, interest in home-based or self-collection options, overall testing experience, and suggestions for improvement (**Figure 3, Supplementary Table 1, Supplementary Figure 2**). Qualitative responses were recorded by the study coordinator, reviewed and analyzed by the principal investigators, and used to refine self-collection training materials and optimize the design of the CD4-FlowSpot.

### 2.6 Flow cytometry

Fresh whole blood samples were stained using the RBC Lysis/Fixation Solution (BioLegend) according to manufacturer instructions (Whole Blood Lyse/No Wash procedure). Cells recovered from dried blood spots were stained according to a proprietary FlowSpot staining protocol. Fresh blood samples were acquired by flow cytometry within four hours of staining; dried samples were acquired within 24-48h after staining. Samples were run and recorded on the 5-laser Cytek Aurora, and unmixing was applied with every acquisition using freshly prepared unmixing controls. The Aurora underwent daily QC by the flow cytometry core staff (Albert Einstein School of Medicine). Antibodies used in this study include CD45 APC, CD15 Pacific Blue, CD14 PE-Cy7, HLA-DR APC-Cy7, CD3 APC-R700, CD4 PE, CD8 PE-Dazzle594, CD19 PE-Cy5, CD3 PE, CD4 BV421, CD45 PerCP, CD69 RB613, IL-2 RB780, TNFα APC, and IFNγ PE. Samples were analyzed using FlowJo (version 10.10.0).

### 2.7 Statistical analysis

Agreement between FlowSpot-derived CD4^+^ T cell percentages and matched venous blood measurements was assessed using multiple complementary statistical approaches. Method comparison was performed using Pearson correlation analysis and Lin’s concordance correlation coefficient (CCC) to evaluate agreement between methods. Deming regression and Passing-Bablok regression were used to assess the constant bias (y-intercept) and proportional bias (slope) across CD4^+^ T cell ranges.

Bland-Altman analysis was performed by plotting the difference between FlowSpot and venous measurements against the mean of the two methods, with mean bias and 95% limits of agreement (LOA) calculated as the mean difference ±1.96 standard deviations. Between-group differences in mean bias were tested with a Welch two-sample *t*-test, and differences in the variability of the paired differences with the Fligner-Killeen test.

Constant and proportional bias were compared by fitting a linear model of FlowSpot on the reference value with a group interaction term (FlowSpot ∼ reference x group), with the group main effect testing for a difference in intercept and the reference-by-group interaction testing for a difference in slope. All tests were two-sided with α = 0.05.

Diagnostic performance for classification of CD4 percentage ≤26% versus >26% was evaluated using a contingency table. Sensitivity was calculated as TP/(TP+FN), specificity as TN/(TN+FP), positive predictive value (PPV) as TP/(TP+FP), and negative predictive value (NPV) as TN/(TN+FN), where TP, TN, FP, and FN represent true positives, true negatives, false positives, and false negatives, respectively. All analyses were performed in R (version 2025.09.2) using paired samples with complete measurements available from both dried finger stick and fresh venous blood specimens.

## 3. Results

### 3.1 Comparison of CD4^+^/CD8^+^ T lymphocytes frequencies in fresh whole blood and FlowSpot across storage conditions

We first investigated the feasibility and stability of flow cytometric immunophenotyping of lymphocytes from blood after drying on a cellulose-based matrix (**Figure 1**). Whole blood was obtained by venipuncture in EDTA from 10 healthy donors. Upon receipt, one aliquot was stained on the same day with a cocktail of fluorescent antibodies targeting T lymphocytes. Additional aliquots from the same donors were applied to five Whatman 903 Protein Saver Cards at 25 µL per spot. Four cards were dried at room temperature (approximately 20-25°C) overnight and then stored in a foil pouch at room temperature (20-25°C) or 37°C for up to 28 days. Samples were then punched with a 6mm hole punch, processed using a proprietary workflow, stained with the same fluorescent antibody cocktail used for fresh samples, and acquired by flow cytometry. The process is summarized in **Figure 1A**. Distinct populations of CD45^+^ leukocytes, CD3^+^ T, CD4^+^ T lymphocytes, and CD8^+^ T lymphocytes were readily identified and quantifiable using FlowSpot, and their percentages were equivalent to that of whole blood processed fresh and on the same day from the same donor for both T cell subpopulations (**Figure 1B**).

**Figure 1.**
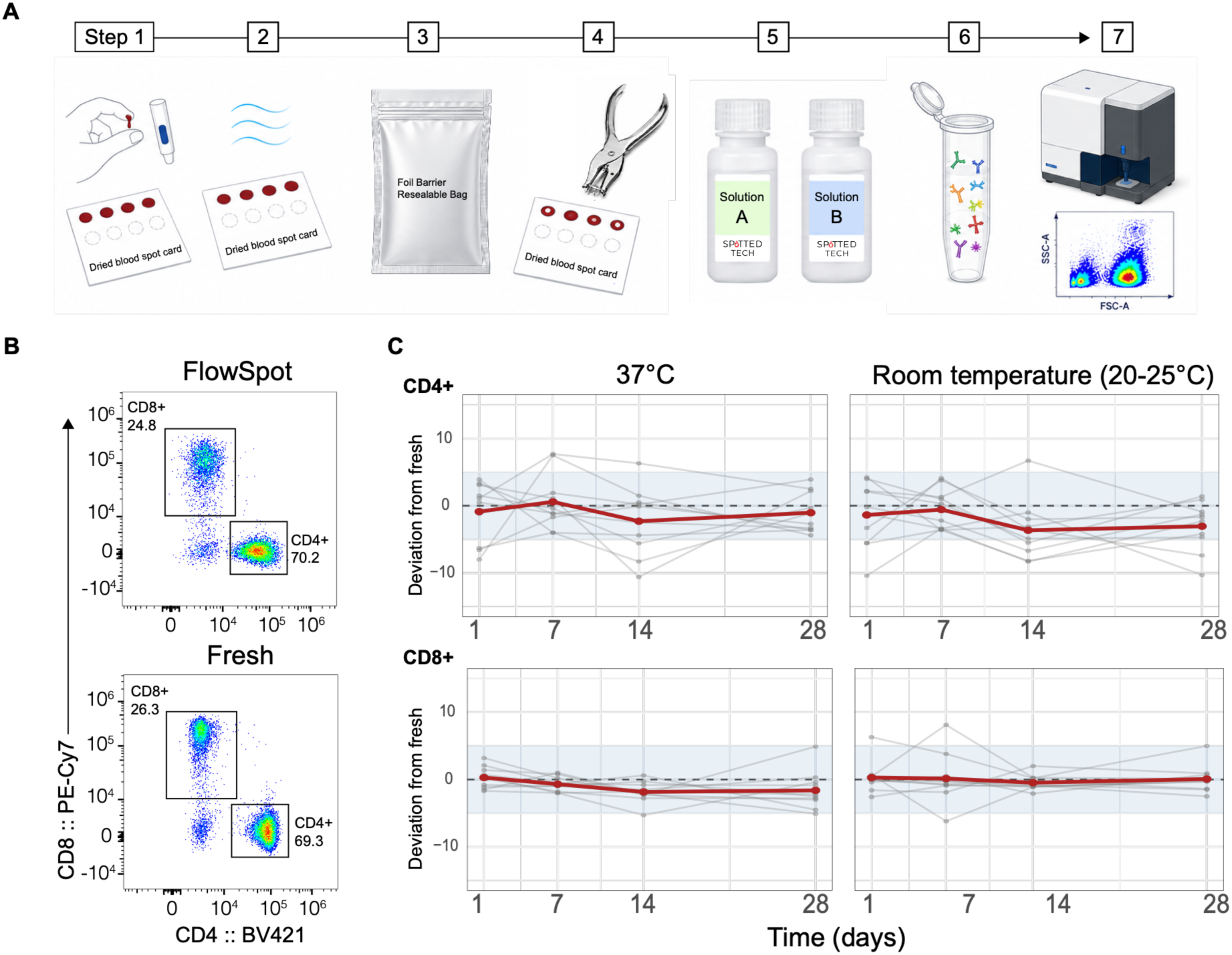
Recovery of T lymphocytes using the FlowSpot system. (**A**) Schematic illustrating the FlowSpot workflow, from blood collection to data analysis. First, blood is obtained by finger stick or venipuncture and dropped onto Whatman 903 Protein Saver Cards. Next, blood is dried and then stored in sealed foil pouches. At time of processing, samples are hole punched with a 6mm punch and processed using proprietary solutions. Recovered cells are stained with a cocktail of fluorescent antibodies and acquired by flow cytometry. (**B**) Representative flow cytometry plots showing CD4⁺ T and CD8⁺ T lymphocytes out of CD3^+^ T lymphocytes recovered with FlowSpot and compared to fresh whole blood stained the same day from the same donor. (**C**) CD4^+^ T and CD8^+^ T lymphocyte deviation analysis of samples from ten healthy donors stored under different conditions (37°C and room temperature) and processed at 1, 7, 14, and 28 days post-collection. Leukocytes are initially gated, followed by lymphocyte gating using CD45^+^ expression and low SSC, single cells are isolated using FSC-A versus FSC-H, followed by identification of CD3^+^ CD4^+^ or CD3^+^ CD8^+^ T lymphocytes out of total singlet lymphocytes. The red line indicates the mean deviation. Grey lines represent individual donors, and the dotted black line at y = 0 indicates no deviation.

We then investigated the effect of storage temperature and duration on the recovery of CD4^+^ and CD8^+^ T lymphocyte percentages using the FlowSpot workflow. Across the 28-day study period, mean bias of CD4^+^ T lymphocyte percentages remained within 5 percentage points of fresh whole blood across storage conditions (**Figure 1C, upper panel**). At 37°C, mean bias relative to fresh whole blood remained within ±2.3 percentage points through 28 days. Room temperature storage produced modest negative bias after 14 days (−3.7 percentage points). The stability of CD8^+^ T lymphocytes as a percentage of lymphocytes was similarly evaluated, and mean bias of CD8^+^ T lymphocyte percentages remained within 3 percentage points of fresh whole blood across all temperatures and time points (**Fig 1C, lower panel**).

Together, these findings demonstrate that the FlowSpot platform preserves accurate T-cell subset quantification for at least 28 days under controlled laboratory storage conditions across a range of temperatures. These results establish the stability of the FlowSpot workflow and provide the foundation for subsequent evaluation under real-world shipping conditions without cold chain requirements.

### 3.2 Clinical validation of the FlowSpot Platform for CD4^+^ T cell enumeration in People Living with HIV using fingerstick collection

Building on the demonstrated laboratory stability of the FlowSpot platform, we next evaluated its performance under real-world shipping conditions. As an initial clinical application, we evaluated CD4^+^ T lymphocyte quantification in people living with HIV (PLWH), enrolling 30 volunteers across two clinics in New Orleans, Louisiana.

Participants provided fingerstick blood samples on Whatman cards at the time of routine, in-clinic visit. In parallel, participants received standard-of-care CD4^+^ T lymphocyte testing from venipuncture blood performed by their clinical providers, and results were shared for comparison with FlowSpot-derived measurements. Deidentified samples were dried, packaged, and shipped by priority or regular USPS mail to the Spotted Tech laboratory in the Bronx, NY, for processing and flow cytometric analysis. Matched reference CD4^+^ T lymphocyte percentages obtained from standard venous blood collected in EDTA and tested at the local clinic reference laboratory were available for 29 participants included in subsequent analysis (**Supplementary Figure 1**). Participant demographics are summarized in **Table 1**. Samples represented a broad range of CD4^+^ T lymphocyte percentages, including values spanning clinically relevant thresholds for HIV monitoring.

**Table 1.** Participant demographics.

| Total (n) | Male (n) | Female (n) | Mean Age<br>(years) | Median Age<br>(years) | Minimum<br>Age | Maximum<br>Age |
| --- | --- | --- | --- | --- | --- | --- |
| 29 | 22 | 7 | 58.2 | 60 | 27 | 85 |

CD4^+^ T lymphocyte percentages measured using the FlowSpot platform showed strong agreement with matched reference measurements obtained by conventional venous blood flow cytometry (**Figure 2**). Passing-Bablok regression analysis demonstrated excellent concordance between methods, with a slope of 1.04 (95% CI: 0.88-1.15) and an intercept of −1.93 (95% CI: −5.62 to 1.99), indicating no significant proportional or constant bias (**Figure 2A**). Similar results were obtained using Deming regression (slope = 1.05, 95% CI: 0.94-1.16; intercept = −1.87, 95% CI: −4.63 to 1.12). Pearson correlation (r = 0.96) and Lin’s concordance correlation coefficient (CCC = 0.960, 95% CI: 0.914-0.980) further demonstrated strong agreement between DBS-derived and reference measurements. Bland-Altman analysis revealed a mean bias of −0.46 percentage points (95% CI: −1.72 to 0.74), which was not significantly different from zero (p = 0.440) (**Figure 2B**). The 95% limits of agreement ranged from −6.71 to +5.78 percentage points, and no significant proportional bias was observed across the measurement range (p = 0.388). Measurement differences were distributed evenly across low and high CD4^+^ T lymphocyte values, supporting consistent assay performance throughout the clinically relevant range.

**Figure 2.**
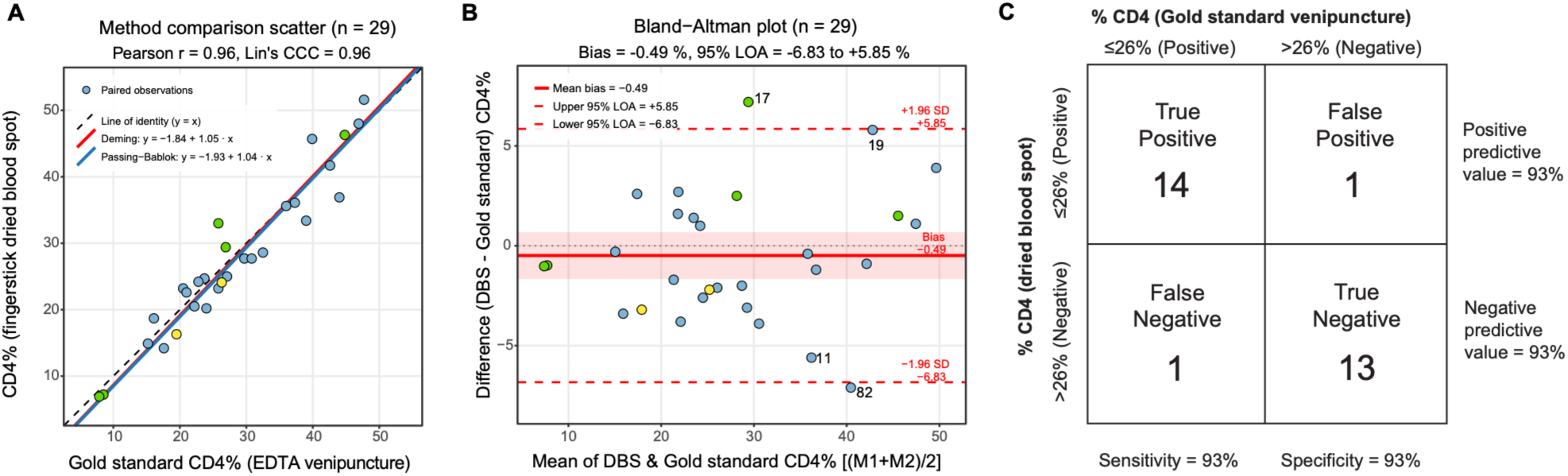
Agreement between FlowSpot-derived and clinically reported percentage CD4⁺ T cells. (**A**) Scatterplot comparing CD4^+^ T lymphocyte percentages out of total lymphocytes using the gold standard clinical data versus FlowSpot. (**B**) Bland-Altman agreement analysis comparing FlowSpot-derived CD4⁺ T cell counts with the gold-standard CD4⁺ measurements, plotted as the difference between the two methods against the gold-standard values. The shaded red region indicates the 95% confidence interval of the mean bias. For (A) and (B), each dot represents one paired observation from one study participant. Yellow dots indicated samples exposed to shipping temperatures that reached 38-45°C. Green dots indicate samples stored for >21 days prior to processing. (**C**) Contingency table evaluating binary classification agreement of CD4^+^ T lymphocytes out of total lymphocytes 26% (considered positive for low CD4^+^ T lymphocytes) or >26% (considered negative for low CD4^+^ T lymphocytes).

To evaluate the utility of FlowSpot-derived CD4% for binary treatment-threshold decisions, we applied a clinically established 26% CD4 cutoff, historically used to guide antiretroviral therapy initiation in resource-limited settings (26), and constructed a contingency table comparing FlowSpot and gold-standard venipuncture classifications across the 29-sample cohort. The cohort was well-suited for this analysis, with 15 of 29 participants (52%) falling below the 26% threshold by venipuncture, providing a near-equal prevalence split that provided balanced power to evaluate both sensitivity and specificity without artificial inflation of either metric by cohort composition. FlowSpot correctly classified 14 of 15 true positive samples (sensitivity 93%) and 13 of 14 true negative samples (specificity 93%), with only one false positive and one false negative across the full cohort. Positive and negative predictive values were both 93% at the observed cohort prevalence. Like the preceding method agreement analyses, the symmetric contingency table indicates that misclassification error was not directionally biased toward either over- or under-estimation of immune compromise by FlowSpot.

### 3.3 FlowSpot performance under standard mail shipping conditions without cold chain requirements

Next, we analyzed the effect of shipment conditions on FlowSpot performance. Variables included time between shipping and processing, temperature exposure, and humidity measurements are shown in **Table 2**. All samples were shipped within three days of collection. Twenty-three samples were processed within 24 hours after arrival and within 14 days of initial collection, and seven samples were stored upon arrival for extended durations before processing. Two samples reached temperatures between 38-45°C during shipping (highlighted in yellow), and seven samples underwent extended storage (in green, **Figures 2A-B**). Both samples that were exposed to higher temperatures, and 5 out of 6 of those that were stored for prolonged time and had matched data, were within 5 percentage points of the gold standard reference data. The full range of time between collection, shipping, and processing ranged from 5 to 153 days, with a mean time between shipping and processing of 34 days. Agreement between FlowSpot and gold standard venipuncture method for CD4 percentages did not differ significantly between the samples that got very hot or underwent extended storage and the remainder of the cohort in constant bias (regression-intercept difference, p = 0.73) or proportional bias (slope difference, p = 0.39), nor in mean bias (p = 0.39) or the spread of the differences (p = 0.92).

**Table 2.** Shipping conditions of PLWH. *Time describes days between shipping and processing.

| Participant | Humidity | Temp (°C) | Time* |
| --- | --- | --- | --- |
| 001-EJ | 30-40 | <38 | 22 |
| 002-EJ | 30-40 | <38 | 13 |
| 003-EJ | 30-40 | NA | 5 |
| 004-EJ | NA | NA | 12 |
| 005-EJ | 30-40 | <38 | 12 |
| 006-EJ | 30-40 | <38 | 12 |
| 007-EJ | 40-50 | 38-43 | 33 |
| 008-EJ | 30-40 | <38 | 15 |
| 009-EJ | 40-50 | <38 | 12 |
| 010-EJ | 40-50 | <38 | 12 |
| 011-EJ | 30-40 | <38 | 12 |
| 012-EJ | 30-40 | <38 | 12 |
| 013-EJ | 30-40 | <38 | 12 |
| 014-EJ | 30-40 | <38 | 12 |
| 015-EJ | 30-40 | <38 | 12 |
| 016-EJ | 30-40 | <38 | 153 |
| 017-EJ | 30-40 | <38 | 113 |
| 018-EJ | 30-40 | <38 | 90 |
| 019-EJ | 30-40 | <38 | 15 |
| 076-TT | 30-40 | <38 | 5 |
| 077-TT | 30-40 | <38 | 5 |
| 078-TT | NA | NA | 12 |
| 079-TT | 40-50 | <38 | 12 |
| 080-TT | 30-40 | <38 | 12 |
| 081-TT | NA | NA | 12 |
| 082-TT | 30-40 | <38 | 12 |
| 083-TT | 30-40 | <38 | 125 |
| 084-TT | 30-40 | <38 | 125 |
| 085-TT | 30-40 | 38-43 | 48 |
| 086-TT | 30-40 | <38 | 34 |

### 3.4 User acceptability of the FlowSpot platform for remote CD4^+^ T cell monitoring in PLWH

A total of 29 participants completed a brief four-question survey immediately after providing a fingerstick blood sample for FlowSpot based CD4^+^ T lymphocyte testing. Overall interest in the technology was high, with a median interest rating of 5 (IQR 4-5). Interest remained consistently strong across age groups, including older adults (≥65 years; n=11), who frequently rated the approach as highly acceptable and emphasized ease and convenience. Younger participants (<40 years; n=5) demonstrated more variable responses (range 1-5) but more often highlighted advantages related to efficiency, time savings, and improved access, particularly for individuals with transportation or work-related constraints.

Reported barriers to conventional blood testing included time off work (n=5), transportation challenges (n=3), needle fear (n=2), psychological burden (n=2), and difficulty obtaining blood (n=1), while 18 participants reported no current challenges. Work-related constraints were more common among younger and middle-aged participants, whereas older participants more frequently reported no major barriers yet still expressed strong support for remote CD4 testing.

Female participants (n=7) uniformly reported high acceptability (ratings 4-5), emphasizing convenience, reduced burden, and improved privacy. Male participants (n=22) similarly demonstrated strong acceptance, with a broader distribution of scores but consistent endorsement of simplicity and practicality of self-collection.

Qualitative feedback highlighted the simplicity, minimal discomfort, convenience, and perceived efficiency of the collection process, with repeated descriptors such as “easy,” “painless,” and “very convenient.” Participants also emphasized benefits for individuals with limited access to care and for enhancing privacy, particularly among younger populations. One participant reported low interest (score 1) despite acknowledging acceptability of the procedure, citing ongoing need for routine clinical management of comorbid conditions. No participants suggested procedural improvements, and several explicitly expressed interest in availability of a home test kit. Collectively, these findings demonstrate strong acceptability of FlowSpot-based CD4^+^ T lymphocyte testing across demographic groups and support its potential as a scalable alternative to clinic-based phlebotomy.

### 3.5 Comprehensive phenotyping on whole blood using FlowSpot

To determine whether additional immune cell types could be identified with FlowSpot, a subset of samples from PLWH were stained with an expanded immunophenotyping panel that enabled identification of CD3^+^ T lymphocytes, CD19^+^ B lymphocytes, CD14^+^ monocytes, and CD15^+^ neutrophils. Each cell type was clearly detected (**Figure 4A**).

**Figure 3.**
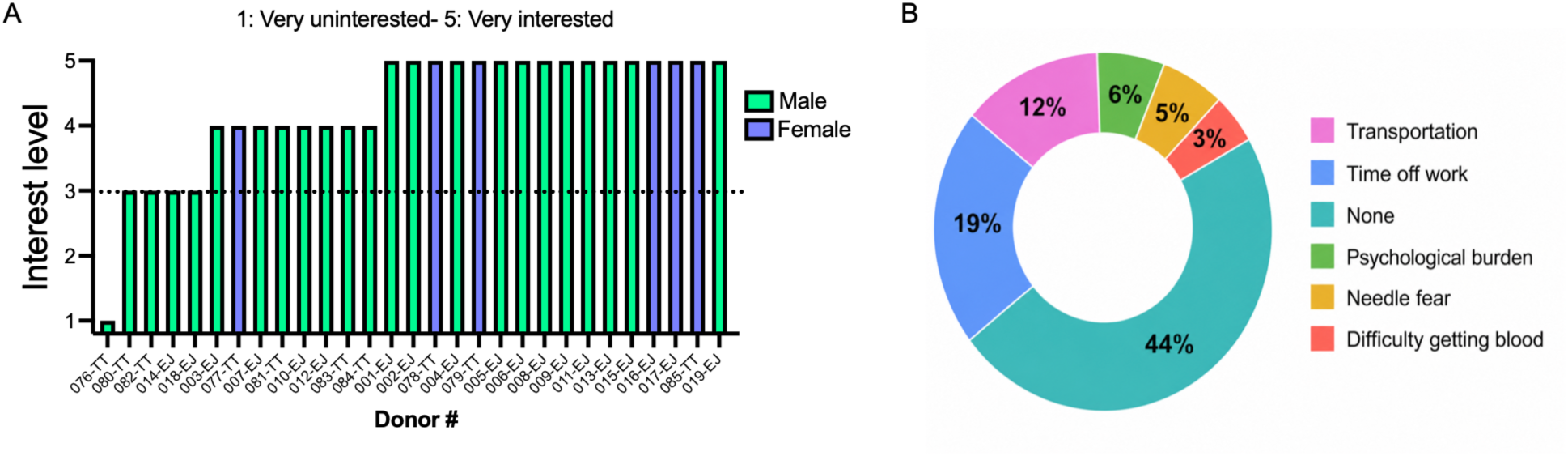
Interest in a home-based CD4-FlowSpot test and current challenges associated with CD4^+^ T lymphocyte monitoring among PLWH. (**A**) Interest levels in a home-based test are shown for individual participants and self-scored on a 5-point scale ranging from 1 (not interested) to 5 (very interested). (**B**) Donut chart summarizing the challenges associated with CD4^+^ T cell level monitoring reported by participants in the questionnaire.

**Figure 4.**
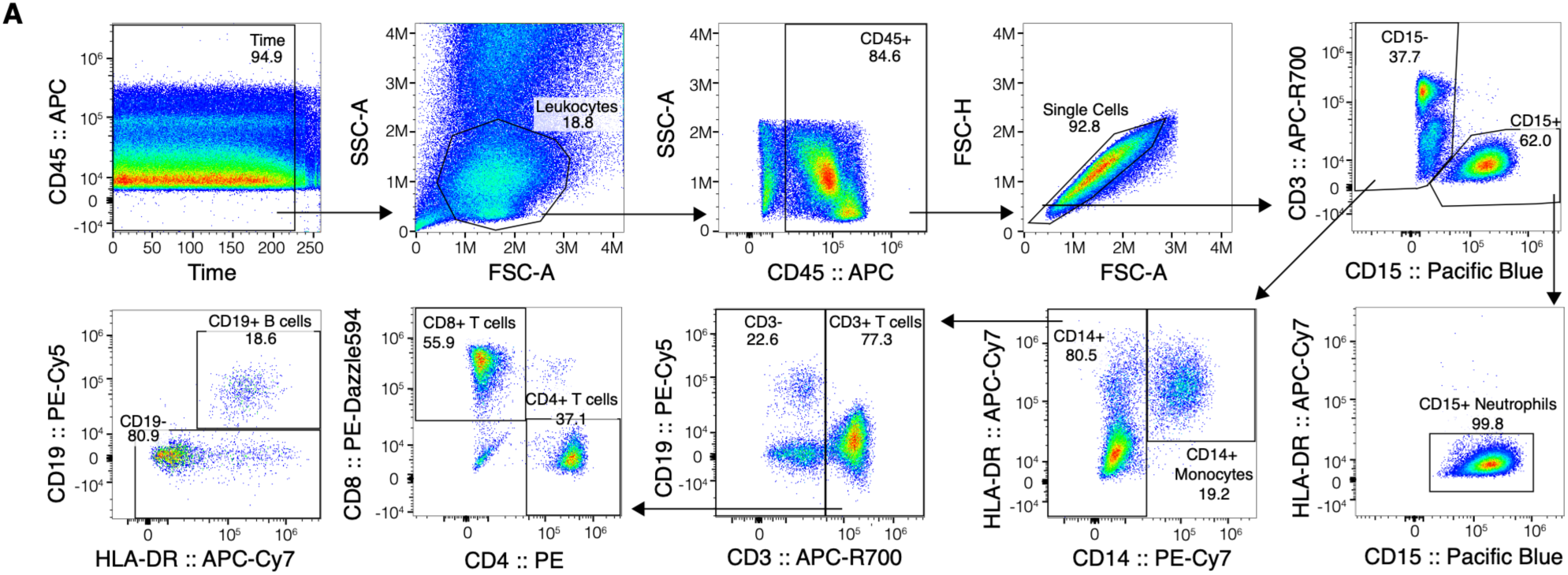
Comprehensive immunophenotyping of whole blood using FlowSpot. (**A**) Representative gating strategy from FlowSpot-derived samples processed 12 days after collection. Samples were first gated on Time vs fluorescence (CD45 APC) to eliminate fluidic artefacts. Next, lymphocytes were gated using FSC/SSC, followed by CD45^+^ pan-leukocytes. FSC-A versus FSC-H were used to identify single cells, and then CD15 was used to separate potential neutrophils from non-neutrophils. CD15^+^ neutrophils were identified by lack of HLA-DR co-expression. From CD15^−−^ cells, CD14 and HLA-DR were used to identify monocytes (CD14^+^HLA-DR^+^) and non-monocytes (CD14^-–^). From CD14-cells, CD3 and CD19 were used to identify CD3^+^ T lymphocytes and CD3-cells. From CD3^+^ T lymphocytes, CD4^+^ and CD8^+^ were used to identify CD4^+^ T helper cells and CD8^+^ T cytotoxic cells. From CD3^-–^ lymphocytes, CD19 and HLA-DR were used to identify B cells (CD19^+^ HLA-DR^+^).

Expected marker co-expression further confirmed accuracy of cell types: CD19^+^ B cells and CD14^+^ monocytes co-expressed HLA-DR, while CD15^+^ neutrophils were HLA-DR negative. Light scatter properties, including forward and side scatter which depict relative size and granularity, differed as expected by cell type: T- and B-lymphocytes had reduced forward and side scatter relative to monocytes and neutrophils (**Supplementary Figure 3**). While paired whole blood data from the same donors were not available, the frequencies of cellular subpopulations fell within expected and previously reported normal ranges.

### 3.6 Intracellular Cytokine Staining using the FlowSpot platform

We next evaluated whether FlowSpot-derived cells retain detectable intracellular cytokines after stimulation. Samples from four healthy donors collected by venipuncture in sodium heparin were stimulated in fresh whole blood for six hours with PMA/ionomycin at 37°C in the presence of protein transport inhibitor Brefeldin A. One aliquot was stained fresh with an intracellular cytokine cocktail containing CD3, CD4, CD8, CD69, IFNγ, TNFα, and IL-2. Additional aliquots of 25 µL were spotted onto DBS cards and allowed to air dry for two hours prior to transferring to a foil pouch. After overnight storage and next-day recovery from DBS, cells were fixed and permeabilized using Cyto-Fast Fix-Perm kit (BioLegend) and stained with the same intracellular cytokine antibody cocktail as fresh. Following stimulation, FlowSpot-derived cells exhibited detectable, moderately decreased expression of IFNγ, TNFα, IL-2, and CD69, with response patterns comparable to those observed in matched fresh blood controls (**Figure 5A-B**).

**Figure 5.**
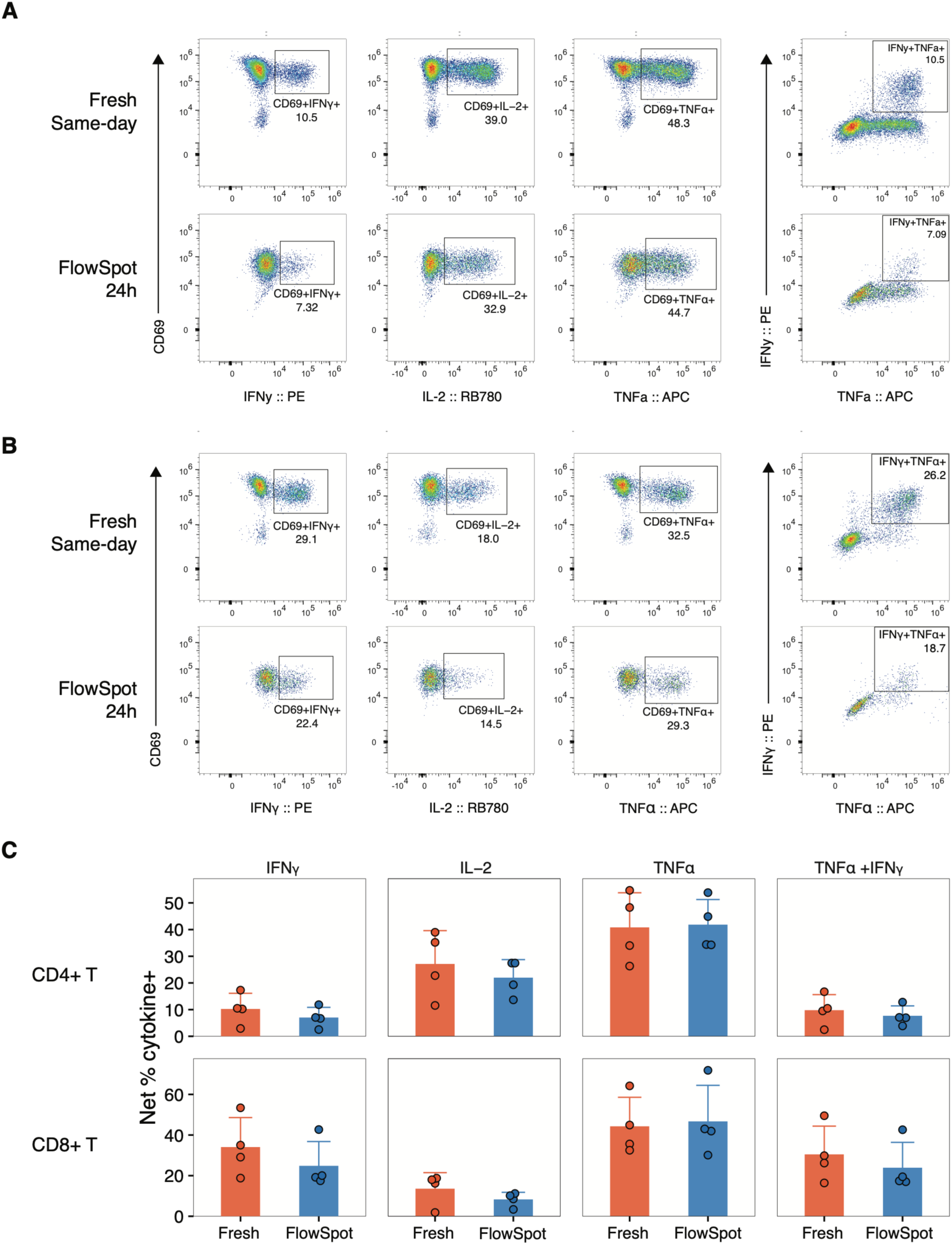
Comparative analysis of intracellular cytokine staining in fresh whole blood versus FlowSpot. Representative flow cytometry plots showing cytokine-producing (**A**) CD4⁺ and (**B**) CD8⁺ T lymphocytes from a healthy donor. Whole blood was stimulated with PMA and ionomycin prior to DBS preparation. For FlowSpot, samples were dried, stored overnight, and processed the following day. Plots show the frequencies of IL-2⁺, IFNγ⁺, TNFα⁺, and cytokine double-positive cells among CD69⁺ CD4⁺ and CD8⁺ T lymphocytes. (**C**) Bar plot comparison of cytokine responses measured in fresh blood and FlowSpot from four healthy donors. Each dot represents an individual donor; bars indicate the mean ± standard deviation. Data points are derived from subtracting the % of events out of stimulated from unstimulated T lymphocytes.

Across donors, frequencies of cytokine-producing cells measured with FlowSpot ranged from approximately 72-106% of those observed in matched fresh samples, depending on the cytokine and T-cell subset (**Figure 5C**). For CD4^+^ T lymphocytes, FlowSpot retained a mean expression of 72% for IFNγ⁺, 88% for IL-2⁺, and 106% for CD4⁺ TNFα⁺ cells, with corresponding mean differences of −3.2, −5.1, and +1.0 percentage points, respectively. Similarly, FlowSpot retained 75% of CD8⁺ IFNγ⁺, 88% of CD8⁺ IL-2⁺, and 105% of CD8⁺ TNF-α⁺ responses, with mean differences of −9.2, −5.3, and +2.4 percentage points, respectively. Despite reductions in response magnitude for some cytokines, FlowSpot-samples retained measurable functional responses across all cytokines evaluated. Importantly, polyfunctional TNFα⁺IFNγ⁺ T lymphocyte responses were also preserved, with FlowSpot measurements averaging 94% of fresh values for CD4⁺ T lymphocytes and 80% for CD8⁺ T lymphocytes.

## 4. Discussion

Blood-based cellular immune testing is central to clinical research, diagnostics, and therapeutic development, providing key insights for biomarker discovery, vaccine evaluation, and immune monitoring. However, it remains heavily dependent on centralized laboratory infrastructure, requiring rapid blood processing, trained personnel, and cold-chain logistics during storage and transport when samples are collected away from testing sites. These logistical constraints limit access to immune monitoring for patients who require frequent testing, such as PLWH undergoing routine CD4^+^ T cell monitoring. On the other hand, they also constrain clinical research by increasing cost and operational complexity, limiting participation in multicenter and longitudinal clinical studies that depend on repeated sampling. As a result, immune monitoring remains a major bottleneck for both routine patient care and precision medicine-driven clinical trials.

Here, we present FlowSpot, a platform designed to overcome these barriers by enabling decentralized, cold-chain-independent immune profiling for clinical diagnostics, at-home monitoring, and clinical research applications. To our knowledge, FlowSpot is the first platform to enable recovery of phenotypically intact leukocytes from conventional cellulose-based dried blood spot cards for downstream flow cytometric analysis.

Notably, it is compatible with commonly used FDA 510(k)-cleared Whatman 903 Protein Saver cards, enabling both T lymphocyte immunophenotyping after prolonged ambient-temperature storage and shipment, and intracellular cytokine staining analysis.

Previous studies have shown that immune cells can be recovered from dried blood under selected conditions (25,27). In 1993, Joseph Yourno reported recovery of leukocytes from blood dried on cellulose-based Guthrie cards, establishing early evidence that cellular components could be preserved in dried specimens. However, this work preceded modern flow cytometric approaches and did not evaluate recovery performance nor compatibility with flow cytometry as the downstream application. More recently, Belkacem and colleagues demonstrated leukocyte recovery from dried blood using a specialized polyester-based matrix (Leukosorb B filter, Cytiva), enabling flow cytometric and transcriptomic analyses and providing important proof-of-concept that dried specimens can retain recoverable cellular information. However, this approach relied on a non-cellulosic proprietary substrate and agitation-based elution methodology, and recovery was not evaluated under decentralized shipping conditions. Unlike polyester, cellulose-based DBS cards are inexpensive are broadly adopted, but retain cells in their fibers, thus presenting greater challenges for leukocyte recovery and compatibility with flow cytometry. FlowSpot retains the accessibility, low cost, and widespread adoption of standard cellulose-based DBS card, while enabling downstream flow cytometric analysis.

Our analytical evaluation demonstrates that FlowSpot preserves quantitative immune measurements with high fidelity. The frequency of T lymphocyte populations, including both CD4⁺ and CD8⁺ T cells, showed high concordance with freshly processed whole blood. The performance of the FlowSpot workflow was maintained following ambient-temperature storage and shipping for up to four weeks, enabling reliable phenotypic immune profiling without cold-chain logistics. This stability substantially expands the practical utility of FlowSpot for decentralized clinical studies, remote patient monitoring, and population-scale immune surveillance.

To rigorously evaluate clinical performance, we tested FlowSpot using fingerstick samples collected from people living with HIV, a population in which regular CD4^+^ T cell monitoring remains clinically relevant even with widespread antiretroviral therapy. We evaluated CD4^+^ T lymphocyte percentages rather than absolute counts, as percentage-based measurements are inherently more robust to variation in blood volume and spot formation during fingerstick self-collection. Compared with matched clinical measurements obtained from fresh blood by venipuncture, FlowSpot achieved approximately 93% sensitivity and specificity, demonstrating that immune measurements obtained from FlowSpot closely approximate conventional laboratory testing. These findings support the analytical validity of the platform while highlighting its potential to extend immune monitoring outside centralized laboratory settings.

Beyond analytical performance, successful implementation of decentralized diagnostics depends on patient acceptance. In our exploratory survey, PLWH reported fingerstick collection as easy, minimally painful, and more convenient than venipuncture, consistent with CDC guidance and prior acceptance of dried blood self-collection (22,28–31).

Participants also highlighted barriers to routine visits, including transportation, time burden, clinic wait times, needle anxiety, and repeated blood draws, which could be alleviated through home-based sampling. Our study was limited to a small clinical cohort at two clinical sites, and samples were collected under supervision following instruction from a study coordinator. Additional studies are needed to assess performance under fully unsupervised at-home collection conditions. Despite these limitations, the ease of capillary blood collection and ambient temperature shipment, combined with strong participant interest in home testing, supports the feasibility of decentralized immune monitoring using the FlowSpot platform.

Intracellular cytokine staining (ICS) by flow cytometry is widely regarded as the gold-standard method for assessing antigen-specific T lymphocyte responses in clinical trials, enabling simultaneous quantification of CD4⁺ and CD8⁺ T cell function and polyfunctionality after antigen stimulation (32–34). ICS is widely used as a correlate of protection in cancer immunotherapy and antiviral vaccine studies, where cytokine-producing T lymphocyte frequencies and functionality have been linked to clinical outcomes in both oncology and infectious disease settings (35–37). However, current ICS workflows are not easily scalable, as they depend on rapid blood processing, viable cell handling, and tightly temperature-controlled transport conditions, limiting their use in large-scale and multicenter clinical studies (34,38,39).

In this study we show that FlowSpot enables ICS-based functional immune profiling. Preserved production of IFNγ, IL-2, and TNFα was detected in CD4⁺CD69⁺ and CD8⁺CD69⁺ T cell subsets, together with comparable Granzyme B expression (data not shown). These findings demonstrate that FlowSpot preserves functional immune readouts in recovered leukocytes in addition to their phenotypic characteristics, suggesting its use in decentralized clinical trials. Although further validation using defined antigen-specific stimulation and systematic evaluation across shipping and storage conditions will be required for clinical implementation, these results establish the feasibility of functional immune profiling from dried blood spots. Moreover, because DBS-based approaches inherently rely on limited sample volumes and antigen-specific T lymphocyte responses occur at lower frequencies than the polyclonal responses evaluated here, further optimization will be required to tailor protocols for these applications. Importantly, all experiments presented were performed using only a portion of a single DBS card, leaving additional sample material available to increase input volume when needed. This remaining capacity provides flexibility to enhance assay sensitivity and support detection of lower-frequency cellular populations, while remaining within the practical limitations of a single DBS card per sample.

Finally, FlowSpot also enabled detection of broader immune populations, including B cells, monocytes, and neutrophils, with frequencies consistent with fresh whole blood, as well as preservation of activation markers such as CD69. However, these markers were not formally evaluated for analytical equivalence and will require further validation. Nonetheless, this capability suggests potential utility for additional at-home diagnostic applications as new blood-based biomarkers emerge, as well as for broader biomarker discovery efforts.

## 5. Conclusion

FlowSpot overcomes a longstanding barrier in cellular immune monitoring by enabling recovery of phenotypically intact leukocytes from conventional cellulose-based dried blood spots, a format historically incompatible with flow cytometric analysis. By removing the need for immediate processing and cold chain logistics while preserving quantitative and functional immune readouts, FlowSpot enables a practical framework for decentralized cellular immune testing. This approach extends immune monitoring beyond specialized laboratories, supporting remote and potentially at-home sampling for both clinical care and research. As healthcare and clinical studies move toward decentralized models, such technologies may enable broader access to immune diagnostics, more inclusive studies, and improved global reach of cellular immune testing.

## Acknowledgments

We thank the study volunteers for their participation, which made this work possible. We also thank the staff at the Albert Einstein College of Medicine Flow Cytometry Core Facility, and we are grateful for the contributions of Sara Babb.

## Author Contributions

M.F. and M.V. designed the study, performed data analysis and interpretation, and provided project oversight. M.V. led preparation of the manuscript. M.F. conceived and led development of FlowSpot technology and methodology, performed experiments, and contributed to the manuscript. R.C. performed experiments, assisted with the development of FlowSpot, and contributed to data collection. S.A. and D.M. coordinated sample collection and shipment, provided fresh whole blood data, and assisted with overall project coordination.

