## Supplementary Files for "FlowSpot Enables Decentralized Phenotypic and Functional Cellular Immune Profiling from Dried Blood Spots"

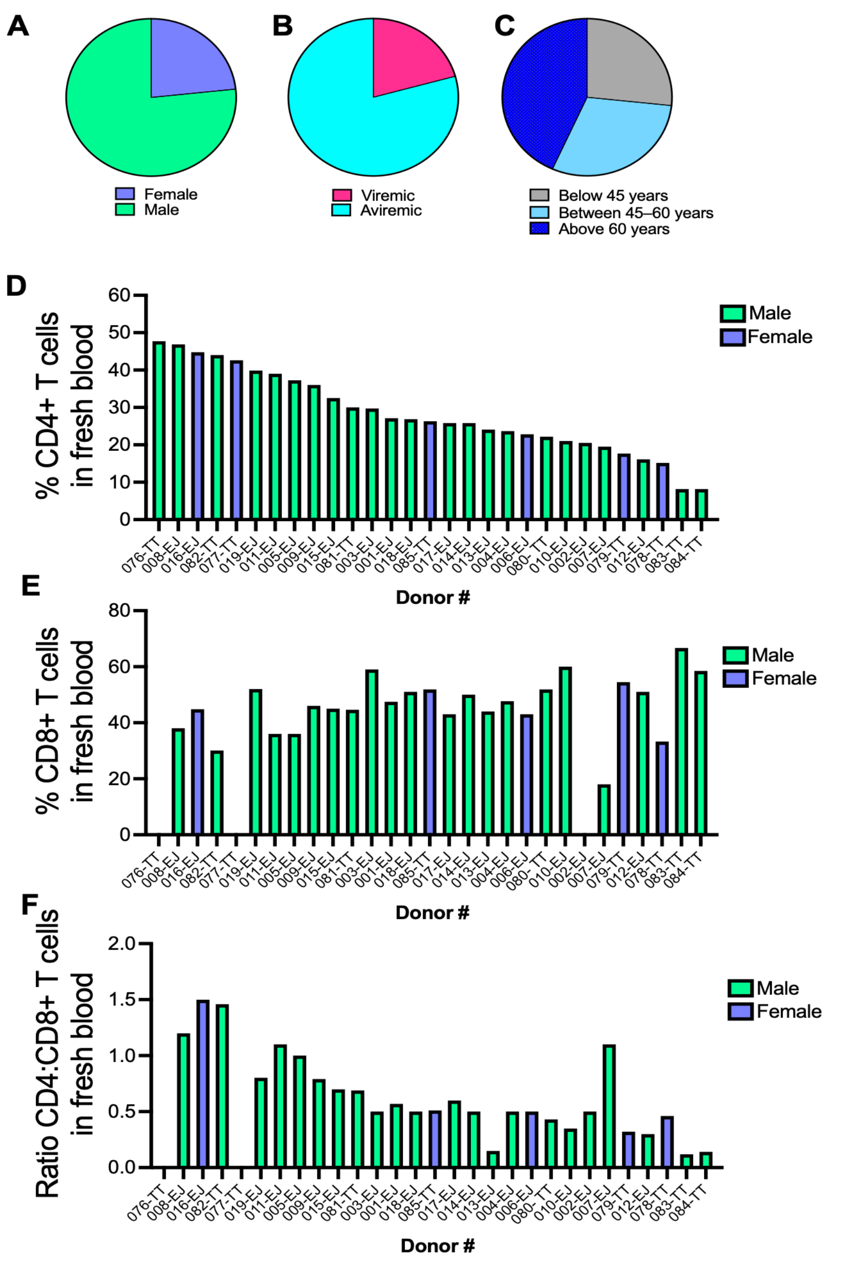


**Supplementary Figure 1. Clinical characteristics and fresh blood T-cell measurements of study participants.**

Pie charts show the distribution of **(A)** sex, **(B)** viral suppression status, and **(C)** age groups among people living with HIV. **(D–F)** Percentages of $CD4^+$ T lymphocytes, $CD8^+$ T lymphocytes, and the CD4:CD8 ratio measured in fresh whole blood. Each bar represents an individual donor and is ordered from highest to lowest $CD4^+$ T lymphocyte percentage across all participants. Clinical and demographic data were obtained from medical records. Blood samples were collected either on the day of the clinical visit or within few weeks before the visit for most donors. Fresh blood T lymphocyte measurements were used as reference values for comparison with dried blood spot-derived measurements.

| PT# | Comments |
|---|---|
| 1 | None |
| 2 | Self-collection is convenient |
| 3 | Simple, painless process |
| 4 | Great idea |
| 5 | Great idea |
| 6 | Love it, easy for patients |
| 7 | Makes things better |
| 8 | Wants home test kit |
| 9 | Self-testing attractive |
| 10 | Good idea |
| 11 | No problem |
| 12 | Good idea |
| 13 | Very easy |
| 14 | Smooth and professional |
| 15 | Convenient |
| 16 | Good idea |
| 17 | Improves privacy and access |
| 18 | None |
| 19 | Good for linkage to care |
| 76 | None |
| 77 | None |
| 78 | Great idea, easy process |
| 79 | Easy for people without transportation |
| 80 | Better access for those without care |
| 81 | Like blood sugar testing |
| 82 | Efficient, saves time |
| 83 | Process fine |
| 84 | None |
| 85 | Yes |

**Supplementary Table 1. Qualitative feedback on CD4-FlowSpot clinic-based testing and perceived acceptability for home use.**

**CD4 Count Testing Questionnaire**

1. **What challenges do you currently face with CD4 count testing?**
   (Please circle all that apply)

   ☐ Transportation
   ☐ Taking time off work for visits
   ☐ Needle fear or pain from venipuncture
   ☐ Difficulty getting blood draws
   ☐ Childcare issues
   ☐ Psychological burden
   ☐ Other (please specify): ____________

2. **How interested are you in self-collecting blood via finger stick and mailing it in to obtain CD4 counts instead of visiting the clinic?**
   (Please select one)

   O 1 – Very uninterested
   O 2 – Uninterested
   O 3 – Neutral
   O 4 – Interested
   O 5 – Very interested

3. **Please provide feedback about the idea of self-collecting blood samples:**

4. **Do you have any suggestions for improving the self-collection process?**

**Supplementary Figure 2: Sample Questionnaire**

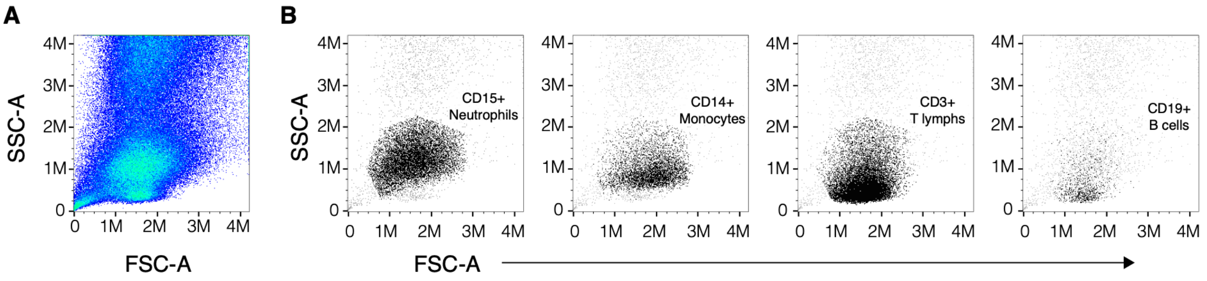


**Supplementary Figure 3. Light scatter resolution with FlowSpot and major cell populations.**

(**A**) Representative forward-scatter-area (FSC-A) and side-scatter-area (SSC-A) light properties representing relative size (FSC-A) and granularity (SSC-A) of events from flow cytometry acquisition. (**B**) Overlay of major cell populations (black) on top of all events to highlight scatter characteristics. An appearance of artificial hard boundaries is a result of gating through the leukocyte gate (**Figure 4A**).
